# Comparing library preparation methods for SARS-CoV-2 multiplex amplicon sequencing on the Illumina MiSeq platform

**DOI:** 10.1101/2020.06.16.154286

**Authors:** Elizabeth M. Batty, Theerarat Kochakarn, Arporn Wangwiwatsin, Khajohn Joonlasak, Angkana T. Huang, Bhakbhoom Panthan, Poramate Jiaranai, Krittikorn Kümpornsin, Namfon Kotanan, Wudtichai Manasatienkij, Treewat Watthanachockchai, Kingkan Rakmanee, Anthony R. Jones, Stefan Fernandez, Insee Sensorn, Somnuek Sungkanuparph, Ekawat Pasomsub, Chonticha Klungthong, Thanat Chookajorn, Wasun Chantratita

**Author notes:** Contributed equally (Elizabeth M. Batty, Theerarat Kochakarn, Arporn Wangwiwatsin). Corresponding author: Elizabeth M Batty.

## Abstract

Genomic surveillance has a key role in tracking the ongoing COVID-19 pandemic, but information on how different sequencing library preparation approaches affect the data produced are lacking. We compared three library preparation methods using both tagmentation (Nextera XT and Nextera Flex) and ligation-based (KAPA HyperPrep) approaches on both positive and negative samples to provide insights into any methodological differences between the methods, and validate their use in SARS-CoV-2 amplicon sequencing. We show that all three library preparation methods allow us to recover near-complete SARS-CoV-2 genomes with identical SNP calls. The Nextera Flex and KAPA library preparation methods gave better coverage than libraries prepared with Nextera XT, which required more reads to call the same number of genomic positions. The KAPA ligation-based approach shows the lowest levels of human contamination, but contaminating reads had no effect on the downstream analysis. We found some examples of library preparation-specific differences in minority variant calling. Overall our data shows that the choice of Illumina library preparation method has minimal effects on consensus base calling and downstream phylogenetic analysis, and suggests that all methods would be suitable for use if specific reagents are difficult to obtain.

## Introduction

Genomic surveillance has played an instrumental role in tracking the Coronavirus Disease 2019 (COVID-19) pandemic, and shed light on the origins and transmission chains in several countries [1–4]. Genomic epidemiology and surveillance are powerful tools for monitoring COVID-19 intervention measures [5, 6]. The use of a multiplex tilling PCR approach developed by the ARTIC network allows the enrichment of virus nucleic acid material for whole-genome sequencing by generating genome-wide amplicons from minuscule amounts of genetic material [7]. This reduces contaminating human genetic material and bypasses virus culture which requires a facility with an advanced biological safety level. The method has successfully been used to amplify and sequence Zika virus from patient samples with Oxford Nanopore Technologies [8], and for multiple real-time genomic surveillance studies on SARS-CoV-2, the etiological virus of COVID-19 [1, 9]. However, limited resources and logistical problems during the time of pandemic has been a bottleneck for many countries to get access to necessary reagents in time, and so availability and price of reagents, and speed of preparation methods, could become major determinants of the sequencing methods used during this crisis. We investigated the effect of three commonly available library preparation approaches (Nextera XT, Nextera Flex, and KAPA HyperPrep) on data quality with multiplex PCR amplicons of SARS-CoV-2 collected in Thailand during the period of March 2020 using the Illumina MiSeq platform. The KAPA HyperPrep method is a ligation-based method, where Illumina-compatible index adapters are directly ligated to amplicon products [10]. The Nextera Flex and Nextera XT kits are two widely-used methods for transposome-based library preparation, where DNA fragmentation and adapter ligation are combined in a single step [11]. The Nextera Flex method additionally uses bead-linked transposomes to allow for easier library normalization [12]. Our method comparison data provides insights into methodological differences on coverage, variant calling and data quality, and could aid decision-making on sequencing approaches in resource-limited settings.

## Methods

### Source of RNA

RNA from SARS-CoV-2 positive patients was acquired from Ramathibodi Hospital, Bangkok after clinical diagnostic tests were performed. All procedures have been approved by Ramathibodi Institutional Review Board (EC approval number: MURA2020/676). C_t_ values for each sample can be found in Supplementary Table 1. C_t_ values for each sample can be found in Supplementary Table 1. Human RNA was used as a negative control. Six SARS-CoV-2 positive samples and two human RNA samples (negative controls) were used for the RT-PCR reaction. All eight samples were then used for preparing three Illumina-compatible library types - Nextera XT, Nextera Flex, and KAPA ligation-based method. All resulting libraries were sequenced on the same run using the Illumina MiSeq platform. The experiment layout is summarised in Figure 1.

**Figure 1.**
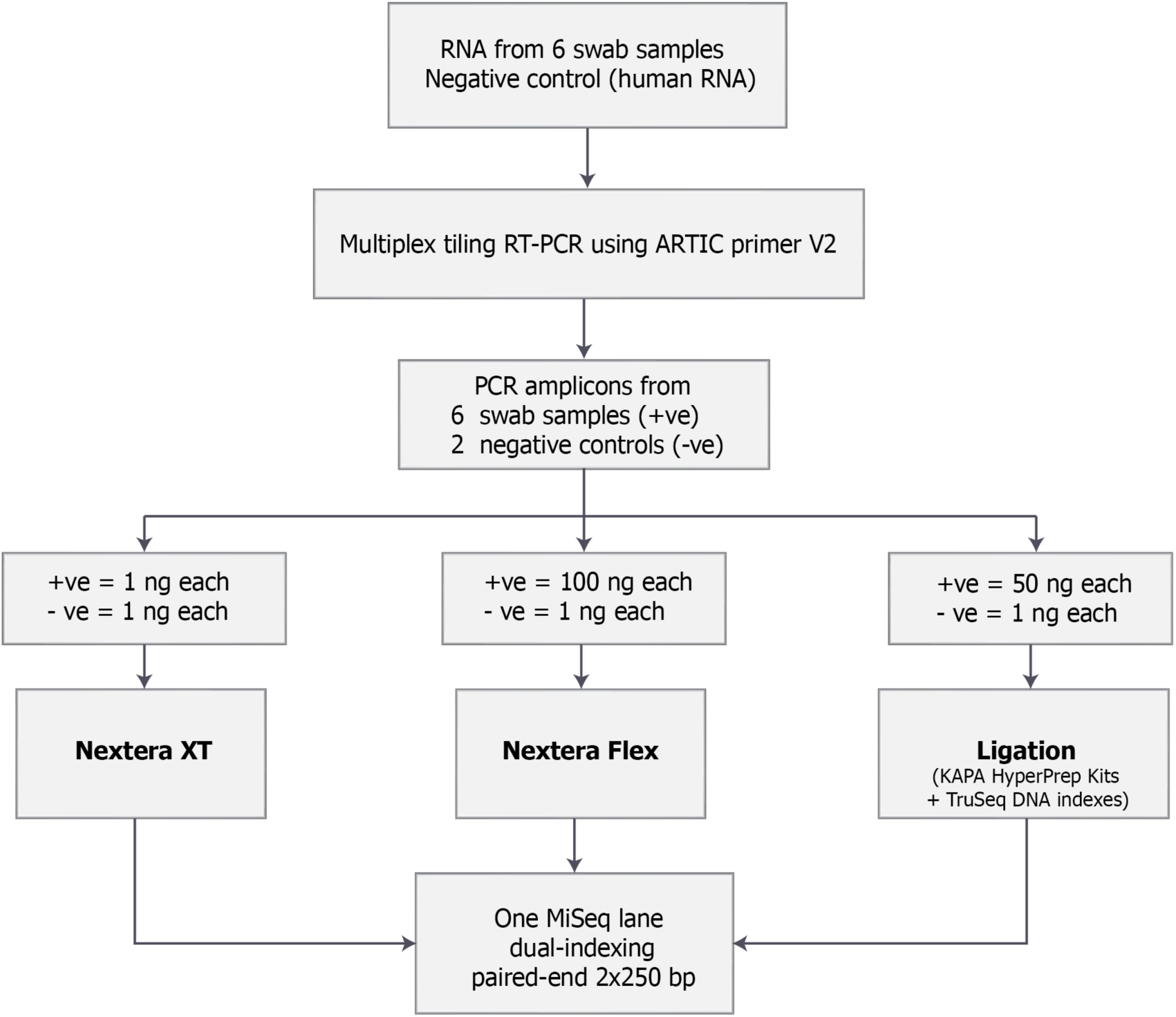
Experimental layout.

### RT-PCR using ARTIC primers

Reverse transcription and multiplex PCR reactions were based on a published protocol from the ARTIC Network [13] using ARTIC primer scheme version 2 which produces ∼400 bp amplicons covering ∼98% of SARS-CoV-2 genome. The primers were provided as two pools, each pool containing 49 pairs. The amplicons from PCR using these primers will overlap each other, but do not overlap within the same pool. Information on the primer sequences can be found at the ARTIC network repository [14]. RNA samples were diluted based on their C_t_ values. Those with C_t_ value between 12-18 were diluted 1:10; those with C_t_ value from 19 upwards were used undiluted. First-strand cDNA synthesis was performed using SuperScript IV reverse transcriptase (Invitrogen, CA, USA). PCR was performed using Platinum SuperFi DNA polymerase (Invitrogen, CA, USA). PCR products were cleaned-up using 1:1 of SPRI beads and eluted in 30µl elution buffer (Qiagen). The resulting amplicons were assayed on the Fragment Analyzer System (Agilent), quantified using Qubit Fluorometer (Qubit dsDNA HS Assay kit, Invitrogen, CA, USA), and diluted to appropriate concentrations for each library protocol.

### Nextera XT library preparation

Amplicons from each pool were diluted to 0.2 ng/µl and 2.5 µl from each pool (0.5 ng) were used as input for each library preparation reaction. For each sample, pool 1 and pool 2 were combined into one library preparation reaction, making a total amplicon input of 1 ng. The library preparation was performed according to Illumina Nextera XT kit protocol. Briefly, the amplicons were incubated with the transposome to perform a tagmentation reaction which fragmented the DNA and added adapter sequences at the 5’ and 3’ ends of each amplicon. The products were then amplified by 12 cycles of PCR using specific index adapters for Illumina sequencing (Nextera XT Index Kit v2, Illumina) (Supplementary Figure 1A). The resulting libraries were cleaned up using AMPure beads (bead:library at 1.8:1 ratio) and eluted in 50 µl of kit resuspension buffer. The negative controls had detectable amounts (>30 ng) after the RT-PCR; therefore, 1 ng was used for Nextera XT library preparation.

### Nextera Flex library preparation

Amplicons from each pool were diluted to 5 ng/µl and 10 µl from each pool (50 ng) were used as input for each library preparation reaction. For each sample, pool 1 and pool 2 were combined into one library preparation reaction, making a total amplicon input of 100 ng. The library preparation was performed according to Illumina Nextera Flex kit protocol. Briefly, the amplicons were incubated with bead-linked transposome to perform a tagmentation reaction which fragmented the DNA and added adapter sequences at the 5’ and 3’ ends of each amplicon. Once tagmented, the fragmented amplicons were cleaned-up and amplified by 5 cycles of PCR using specific index adapters for Illumina sequencing (Nextera™ DNA CD Indexes, Illumina) (Supplementary Figure 1B). The resulting libraries were cleaned up using AMPure bead (bead:library at 1.8:1 ratio) and eluted in 30 µl of kit resuspension buffer. For the negative controls, 1 ng was used for Nextera Flex library preparation.

### KAPA ligation-based library preparation

Amplicons from each pool were diluted to 1 ng/µl and 25 µl from each pool (25 ng) were used as input for each library preparation reaction. For each sample, pool 1 and pool 2 were combined into one library preparation reaction, making a total input amplicons of 50 ng. The library preparation used KAPA Hyper Prep kit (Roche), and the protocol was based on a published method [7] and the kit manual using full reaction volume. Briefly, the amplicons were end-repaired and A-overhang added. The A-overhanged products were ligated with index adapters for Illumina sequencing (TruSeq DNA CD Indexes, Illumina). Post-ligation products were cleaned up using AMPure beads (bead:library at 0.8:1 ratio) and eluted in 25 µl. Then, 20 µl were used for library amplification by 8 cycles of PCR using KAPA Library Amplification Primer Mix (Supplementary Figure 1C). The resulting libraries were cleaned up using AMPure bead (bead:library at 1:1 ratio) and eluted in 30 µl using nuclease-free water. For the negative controls, 1 ng was used for ligation-based library preparation.

### Library QC

All libraries were assayed on Fragment Analyzer System (Agilent) and quantified using Qubit Fluorometer (Qubit dsDNA HS Assay kit, Invitrogen) prior to sequencing. For Nextera XT and Nextera Flex, the fragment sizes of amplicon insert plus sequencing adapters were 259-320 and 440-470bp, respectively. For the KAPA ligation-based method, fragment sizes were 512-556bp (Supplementary Table 2).

### Library pooling and sequencing

All libraries were diluted to 2 nM. Same volume of each 2 nM library was taken to produce a equimolar pool of 24 libraries. Of the pool, 5 µl was mixed and incubated for 5 minutes at room temperature with 5 µl 0.1 N NaOH to denature the dsDNA. PhiX was also denatured using 0.1 N NaOH. The denatured library pool and PhiX were each diluted to 10 pM by adding HT1 solution from the MiSeq kit. The library pool, HT1 solution, and 5 µl 20 pM PhiX were mixed to achieve 8 pM final library concentration at a total volume of 600 µl. The pool was sequenced on one MiSeq run using MiSeq v2 chemistry with 250bp paired-end sequencing.

### Sequencing data QC

To identify human contamination, reads were mapped to both the human GrCH38.p11 and the SARS-CoV-2 genome MN908947.3 using Bowtie2 [15]. All reads which mapped to the SARS-CoV-2 genome were uploaded to the Sequence Read Archive under project PRJEB38369.

Variant calling was performed with the ncov2019-artic-nf pipeline with default Illumina parameters (commit f0ba0a1493c9571f4eda161ae8d6afe02d0da570) [16]. In this pipeline, reads were mapped to the MN908947.3 reference genome using bwa 0.7.17-r1188 [17]. TrimGalore 0.6.5 [18] was used to trim reads for adaptors and low-quality regions and Ivar v1.1_beta [19] was used to trim primer regions, call variants, identify primer mismatches and generate consensus sequences. Reads that did not begin with a primer were retained using the ‘-e’ option for the two Nextera library preparation methods but not for the KAPA libraries. Variants in primer regions were identified by Ivar getmasked. Ivar variants was run with the parameters minimum quality score 20, minimum threshold 0.1, and minimum coverage 50 to identify minor variants.

Reads were downsampled using seqtk 1.3-r106 [20] and run through the same pipeline to call variants and generate consensus sequences. Coverage per amplicon was determined using bedtools v2.29.2 [21].

## Results

RT-PCR using ARTIC primers generated detectable concentration for all samples. The negative controls yielded some product, but the amounts were lower than that of the SARS-CoV-2 positive samples by at least 10-fold. No correlation of C_t_ values with amplicon yields could be seen in this small sample. The PCR yield from pool 1 and pool 2 was concordant. For library preparation, the smallest to largest insert size was Nextera XT, Nextera Flex, and KAPA ligation-based method, respectively The highest library yield was from the KAPA ligation-based method, followed by Nextera XT, and Nextera Flex. All three methods yielded enough library product for use in the sequencing (Supplementary Table 2).

All 18 libraries were run together on a single MiSeq run. 812,984 to 1,662,778 sequence reads were generated for each positive sample library. The number of reads generated and the percentage of reads which could be mapped to the SARS-CoV-2 reference genome were similar across library types (Figure 2A and 2B).

**Figure 2.**
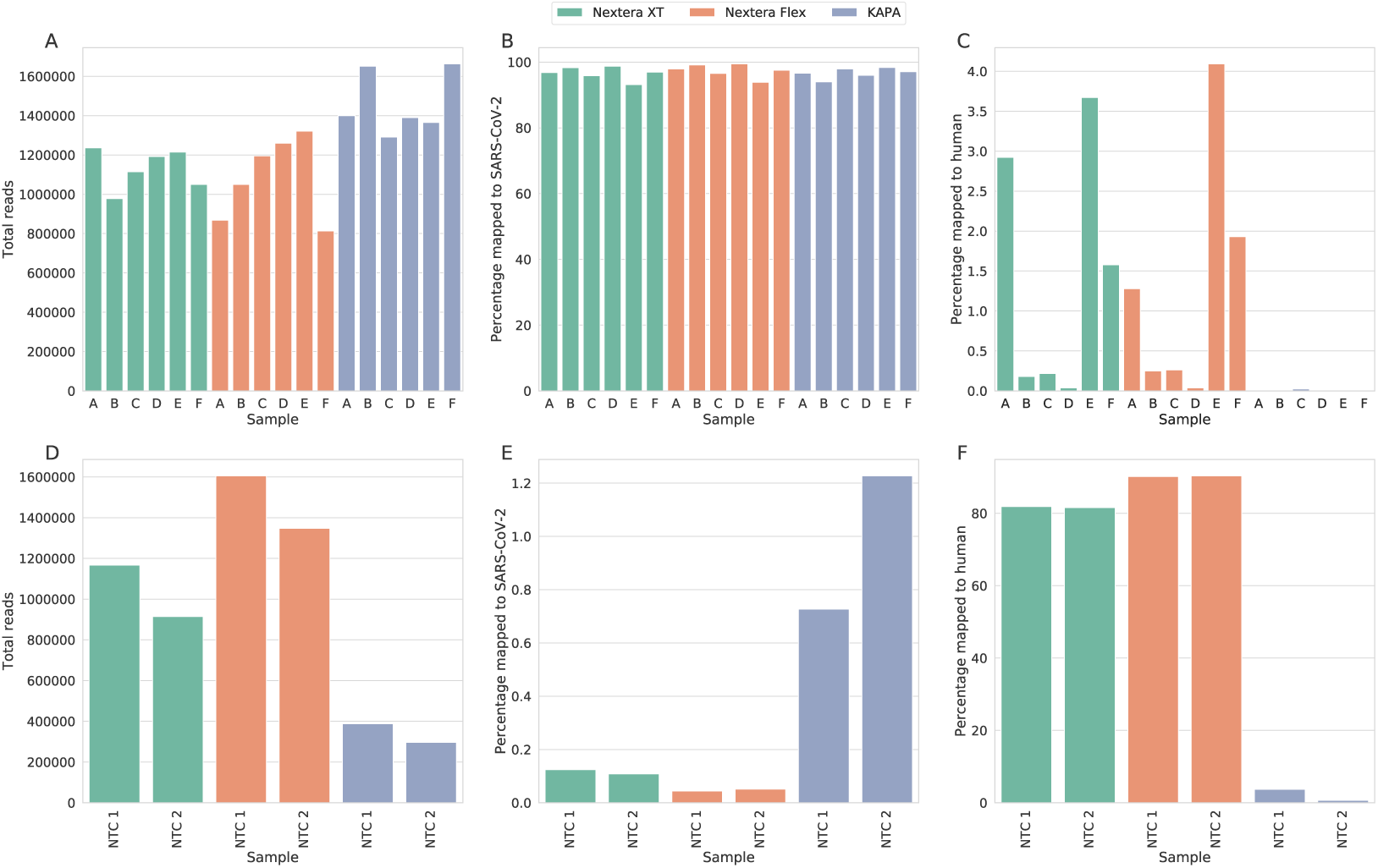
Total number of reads and percentages of reads mapped to SARS-CoV-2 or human reference genomes. Top row shows A) total reads, B) percentage mapped to SARS-CoV-2 reference genome, and C) percentage mapped to the human reference genome for six samples prepared using the three different library prep methods. The bottom row shows D) total reads, E) percentage mapped to SARS-CoV-2 reference genome, and F) percentage mapped to the human reference genome for two negative control (NTC) samples prepared with the three different library prep methods.

By mapping the reads to the human genome we could assess levels of human DNA contamination in each sample. Human contamination was lower in the KAPA-prepared samples (0.01-0.02% of reads mapped) than in the Nextera-prepared samples (0.03-3.67% in XT and 0.04-4.09% in Flex) (Figure 2C).

We included two negative control samples containing human RNA which were taken through the complete PCR and sequencing process alongside the positive samples. More reads were generated using the Nextera methods than the KAPA library preparation from these negative control samples. The negative control samples prepared using the Nextera methods had high levels of human sequences (74.21-83.3%) while the negative controls prepared with KAPA showed low levels of human sequences (0.57% and 3.29%) (Figure 2D). A low percentage of reads can be mapped to SARS-CoV-2 (Figure 2E), and after the filtering and primer trimming steps of our pipeline many of the reads were removed, suggesting they were low-quality or (in the case of the KAPA ligation-mediated preparation) did not start with primer sequences (Supplementary Table 3).

### Genome and amplicon coverage

As we use amplicons as our input material for sequencing, the coverage across the genome will be influenced by the efficiency of the amplicon PCR. Figure 3 shows the median normalised coverage for each amplicon, showing there was high variability between amplicons. This data was produced using the ARTIC V2 amplicon set, which has known problems with amplicons 24, 54, 64, 70, and 74 [22].

**Figure 3.**
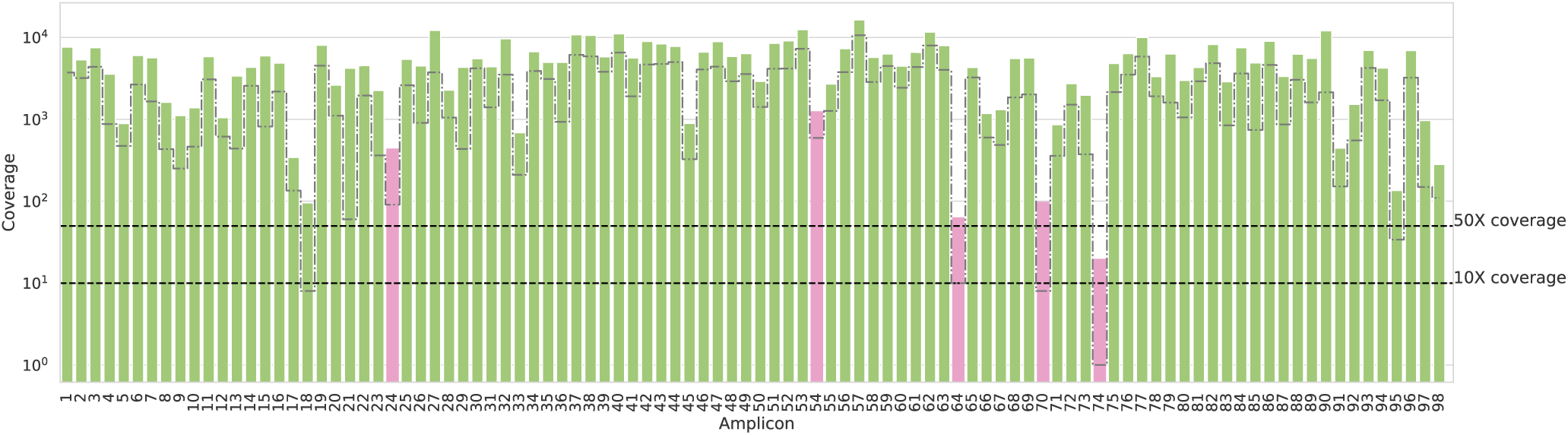
Plot of median coverage for each amplicon across all positive samples. Dotted lines mark 50x coverage (our target coverage level) and 10x coverage (minimum depth needed to call variants in our pipeline). The dot-dash line marks the lower bound of coverage in our samples. The bars plotted in pink are those which are known to have poor amplification in the ARTIC v2 primer set.

To determine if there were differences in amplicon coverage between the different library preparation methods using the same input PCR, we compared the coverage of all amplicons for each library prep method (normalising the data within each library). The plot suggests a slight difference in the coverage between the Nextera and the KAPA library prep methods, with the KAPA-prepared amplicons showing more amplicons with high coverage, but the difference was not statistically significant (using the Mann–Whitney–Wilcoxon test with Bonferroni correction for multiple testing) (Figure 4).

**Figure 4.**
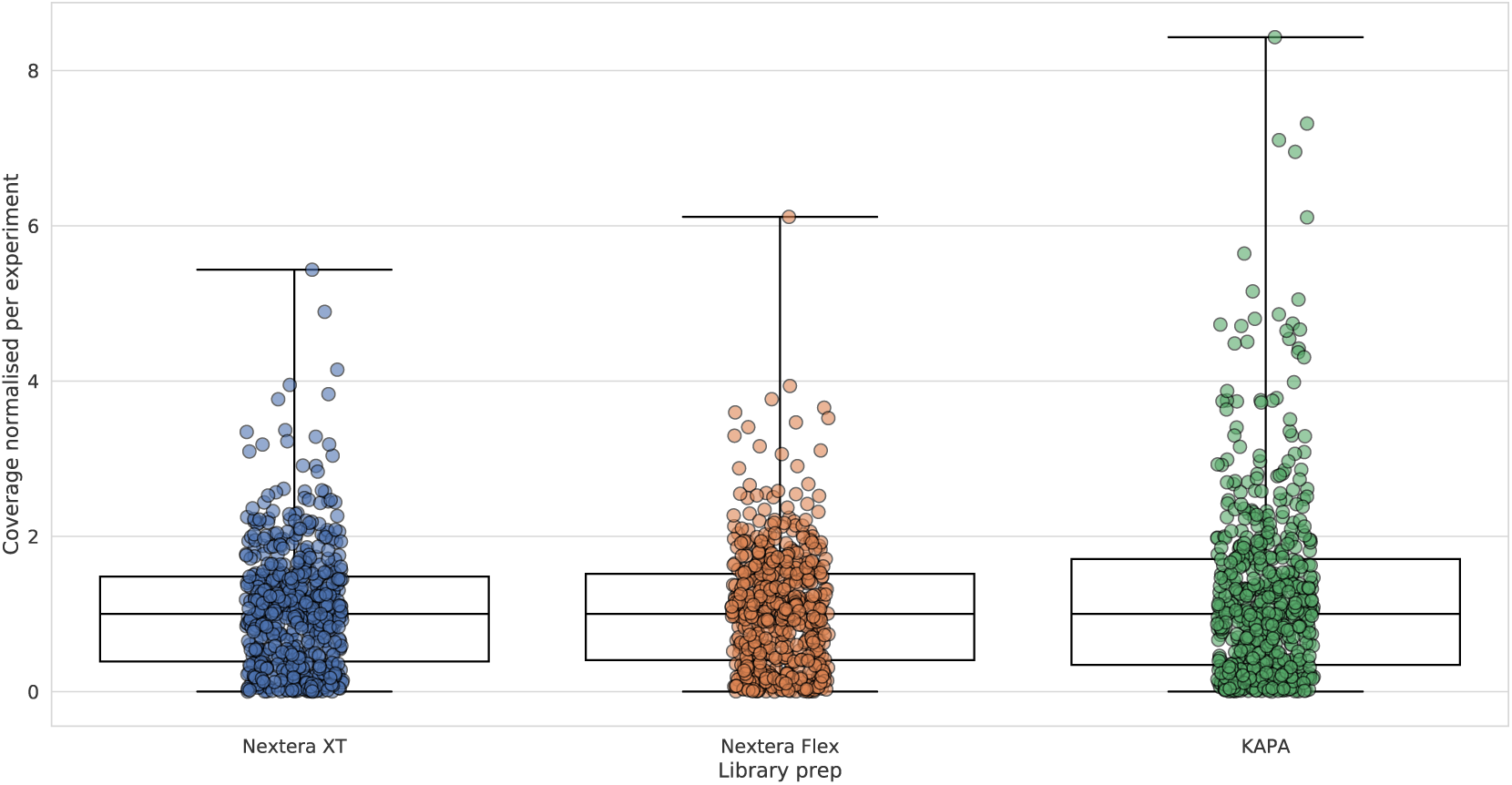
Coverage across all amplicons for each library prep method. Coverage was normalized for each experiment, and all amplicons from across the 6 positive samples are plotted for each library preparation method.

### SNP calling

To see how library preparation method and coverage affect SNP calling, we first used all reads to call SNPs against the reference genome. For each sample, the consensus sequences generated from different library preparations gave identical SNPs relative to the reference genome (6-9 SNPs depending on sample). Samples A and F had SNPs called in the primer region for amplicon 97, and do show lower coverage at this amplicon across all library preparation methods compared to other samples (Supplementary Figure 2).

### Downsampled data

To determine how much sequence data is required, we downsampled the reads for each sample to the same level and tested a range of read pairs between 400,000 and 10,000 to determine the minimum number of reads needed to cover the whole genome and call all SNPs in these samples.

Samples with 250,000 read pairs and above gave identical results to the full read sets when calling SNPs against a reference. With 100,000 read pairs, a cluster of 3 SNPs at positions 28881-28883 in sample A was missed using the Nextera XT prep; however in all other cases 100,000 read pairs was sufficient to call all SNPs (Supplementary Figure 3).

We looked at the proportion of the genome where coverage falls below the levels needed for variant calling. Figure 5 shows the number of bases which fall below 10X and 50X coverage for different downsampled read sets. Different completeness thresholds have been used for different analyses - for instance, the Nextstrain platform [23] will filter out samples with <27,000bp called [24], while GISAID [25] will mark as complete those strains with <29,000bp called. We used these two thresholds to assess our data. With 10X coverage required, the 27,000bp threshold was achieved by as few as 25000 read pairs, while 29,000bp was reached using 250,000 read pairs in all libraries prepared by the Nextera Flex or KAPA methods, and 4 out of 6 libraries prepared using the Nextera XT library prep method. The 29,000bp threshold was not consistently reached at 50X coverage for any set of data, likely due to amplicon dropout; the 27,000bp threshold was covered at 50X using 100,000 read pairs in all libraries prepared by Nextera Flex or KAPA methods, and 5 out of 6 libraries prepared using the Nextera XT method.

**Figure 5.**
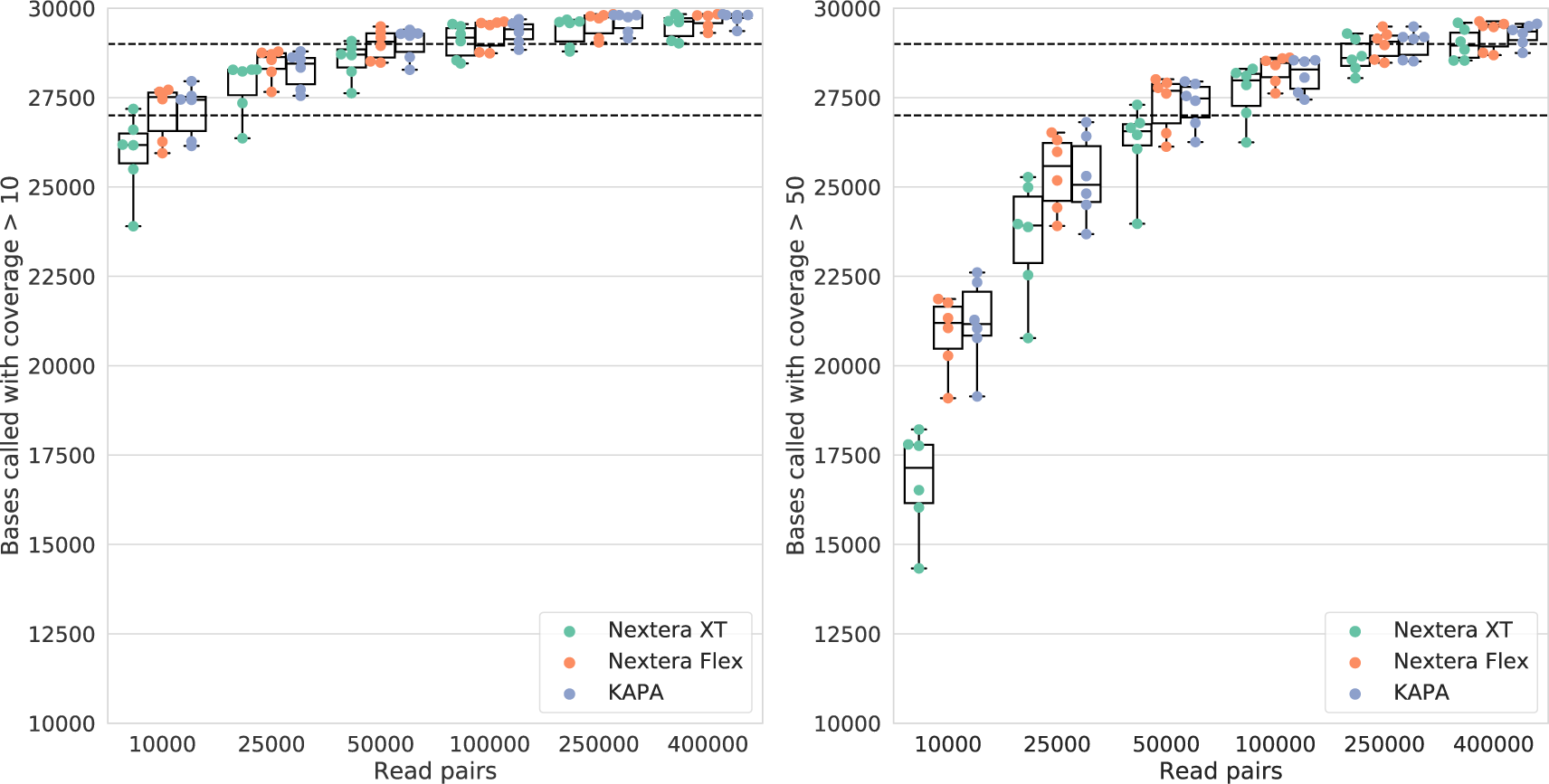
Effects of downsamping reads on consensus base calling. Panels a) and b) show the percentage of the genome which is above the threshold of a) 10X coverage and b) 50X coverage, coloured by library prep method. Horizontal dashed lines mark quality thresholds of Nextstrain (27,000bp) and GISAID (29,000bp).

When we compare the three different library prep methods, at high numbers of read pairs the results are comparable, however the KAPA and Nextera Flex methods give better genome coverage at low numbers of read pairs than the Nextera XT method.

While we are primarily concerned with reporting the consensus variants in these samples, amplicon sequencing is capable of reporting intrahost diversity [19]. We examined our variant calls for evidence of intrahost diversity to determine if this was affected by library prep methods. We identified all variants called with a frequency between 0.10 and 0.99 using a minimum depth of 50 and compared whether they were called across methods and with the same frequency. A total of 24 positions were called as minority variants in one of six samples, with the majority of variants falling around 10% frequency (Figure 6). Noticeably, the same position was called as a minority variant in 5 of our 6 samples using the KAPA library prep method, despite the samples showing few consensus shared variants. On inspection of the alignment, the non-reference base calls were found only in negative strand reads. This variant fell into a region which was covered only by negative strand reads in the KAPA library prep, while in the Nextera preps there were reads from both strands and the ratio of reads with this variant is lower (Supplementary Figure 4).

**Figure 6.**
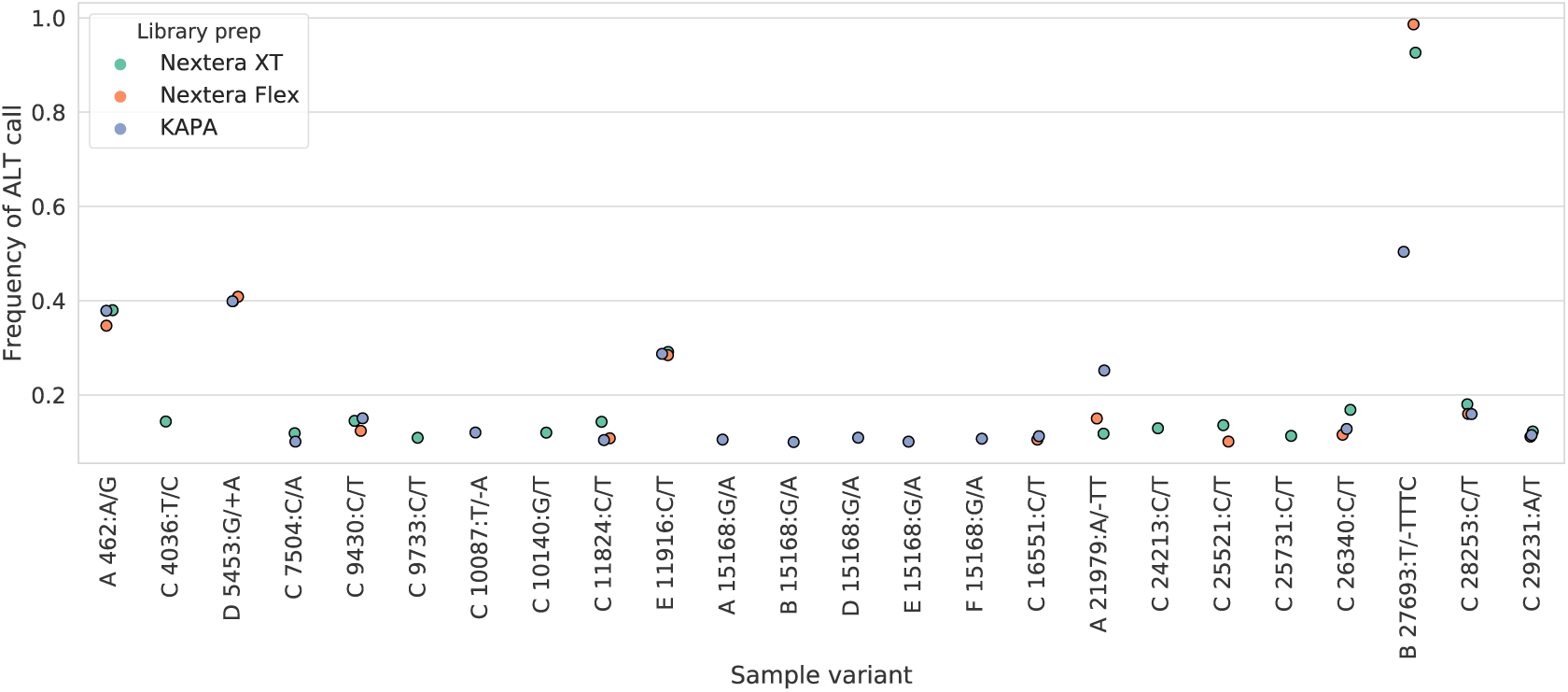
The frequency of the ALT base in variant calls across the six samples, coloured by library prep method. Only variants where the ALT frequency is >0.1 and < 0.99 are included.

Of the four variants which were found at >20% frequency by all different library prep methods, three show good concordance between the three methods. However, one variant, a 4bp deletion at position 27693 which was validated by Sanger sequencing[26], is found at a high frequency in the Nextera library prep samples and at a lower frequency in the KAPA library prep sample. On inspection of the alignment, this is due to the variant falling in a homopolymer tract which is 247bp from the start of amplicon 91 in our scheme. In the KAPA library, as the indel falls near the end of the read on the positive strand, the bases are soft-clipped in the alignment and do not call the deletion, giving a frequency of 50% as only the negative strand reads show the deletion (Supplementary Figure 5). As the reads in the Nextera libraries have varying start positions, only a small number are soft-clipped and this does not have a large effect on the variant frequency.

## Discussion

By comparing the same SARS-CoV-2 samples across different library prep methods, we can assess the effectiveness of the different methods to get complete genome coverage and SNP calling. We conclude that all three methods give acceptable data, and at high coverage levels the regions which cannot be called due to low coverage are more affected by the success of the amplicon PCR and not by the library prep methods. Using a conservative cutoff of 50X coverage at all positions, we could call over 95% of the genome using all methods, and the same SNP calls were seen across all library prep methods, indicating that the method of preparation should not affect base calls or downstream phylogenetic analysis. We saw higher human DNA contamination in the Nextera library preparations, however this did not affect the downstream analysis.

By including negative control samples alongside our positive samples, we could see that the negative controls from the transposase-based Nextera methods and the ligation-based KAPA method behave differently. While the transposase-based methods have a higher number of reads in the negative controls, more of these reads are human contamination. A small number of reads remain after the filtering and trimming stage, indicating that few high quality SARS-CoV-2 reads were retained in these samples, and the small numbers remaining may be due to index hopping from the positive samples. This suggests that the recommendations of the ARTIC network be followed and unique dual indexes or unique molecular indexes used for barcoding, along with including a negative and positive control sample on each run [27].

Our PCR amplicons were generated with the ARTIC V2 primer set, which has known amplicons which consistently drop out, and has already been replaced with an updated version which should reduce these problems. While we saw amplicon drop-out in agreement with those previously reported for this primer set in amplicons 64,70, and 74, we also saw low coverage in amplicon 18, which was previously noted to have low coverage in ARTIC V1 due to primer dimer formation but not in V2. The presence of SNPs in the primer region for amplicon 97 in two of our samples appears to affect PCR efficiency for this amplicon. While our results are broadly in line with those previously reported, the unexpected dropout of amplicon 18 suggests that amplicon efficiency should be further monitored, and locally circulating sequence variants in primer regions may affect amplification of those amplicons.

We downsampled our data to determine a minimum level of coverage and read pairs needed to generate usable data. The Nextera Flex and KAPA prep methods produced very similar results in terms of SNP calling and genome coverage. The Nextera XT prep required a higher number of read pairs to produce the same level of coverage.

250,000 read pairs using 250bp paired end reads was sufficient to call over 29,000bp of the genome at 10X coverage with the Nextera Flex and KAPA methods. Assuming 24 million read pairs are produced from a single MiSeq run using the MiSeq v2 reagent kit [28], this suggests that up to 96 samples could be pooled on a run. Using the V3 amplicon PCR primers should improve the coverage figures and potentially allow for fewer reads to be used. A lower threshold of calling 27,000bp would be reached by all of the Nextera Flex and KAPA samples with only 100,000 read pairs per sample, and allow for a high number of samples to be multiplexed on a single run by trading complete genome coverage for reduced sequencing costs. However, as library preparation costs become a larger proportion of the cost per sample, the savings may be small.

While we have few samples to investigate minority variant calls, our limited data shows that while there is good agreement between different library preparation methods on variants present at above 20% frequency, there are method-specific differences which are largely due to differences in the positive and negative strand reads. Due to the nature of amplicon sequencing, it is not possible to require reads from both strands to be confident of the presence of a variant, and it is recommended to carefully filter and further investigate any minority variants which appear to be present in reads from one strand only or are amplified in only one amplicon.

## Declarations

### Ethics approval and consent to participate

All procedures have been approved by Ramathibodi Institutional Review Board (EC approval number: MURA2020/676). RNA samples were surplus material from clinical diagnostic testing.

### Availability of data and materials

The raw sequence reads which map to SARS-CoV-2 are available from the ENA as project PRJEB38369.

### Authors’ contributions

EMB, TK, AW and TC prepared the original draft. EMB, TK, AW, AH, NK performed data analysis and visualisation. AW, KJ, BP, KK, WM, TW, KR, IS, SS and EP performed the experiments. ARJ, ST, CK, TC and WC supervised and administered the research. TC and WC acquired funding for the project. All authors reviewed and edited the final article.

## Acknowledgements

The computational aspects of this research were supported by the Wellcome Trust Core Award Grant Number 203141/Z/16/Z and the NIHR Oxford BRC. The views expressed are those of the author(s) and not necessarily those of the NHS, the NIHR or the Department of Health. EMB is supported by a Wellcome Trust Core Grant to the Thailand Major Overseas Programme (090532/Z/09/Z). Funding was also received from the Thailand Center of Excellence for Life Sciences and National Research Council of Thailand. Material has been reviewed by the Walter Reed Army Institute of Research. There is no objection to its presentation and/or publication. The opinions or assertions contained herein are the private views of the author, and are not to be construed as official, or as reflecting true views of the Department of the Army or the Department of Defense.

## References

1. Lu J, du Plessis L, Liu Z, Hill V, Kang M, Lin H, et al. Genomic epidemiology of SARS-CoV-2 in Guangdong Province, China. medRxiv. 2020;:2020.04.01.20047076. doi: 10.1101/2020.04.01.20047076.

2. Oude Munnink BB, Nieuwenhuijse DF, Stein M, O’Toole Á, Haverkate M, Mollers M, et al. Rapid SARS-CoV-2 whole genome sequencing for informed public health decision making in the Netherlands. bioRxiv. 2020;:2020.04.21.050633. doi: 10.1101/2020.04.21.050633.

3. Bedford T, Greninger AL, Roychoudhury P, Starita LM, Famulare M, Huang M-L, et al. Cryptic transmission of SARS-CoV-2 in Washington State. medRxiv. 2020;:2020.04.02.20051417. doi: 10.1101/2020.04.02.20051417.

4. Seemann T, Lane C, Sherry N, Duchene S, da Silva AG, Caly L, et al. Tracking the COVID-19 pandemic in Australia using genomics. medRxiv. 2020;:2020.05.12.20099929. doi: 10.1101/2020.05.12.20099929.

5. Kümpornsin K, Kochakarn T, Chookajorn T. The resistome and genomic reconnaissance in the age of malaria elimination. Dis Model Mech. 2019;12. doi: 10.1242/dmm.040717.

6. Gardy JL, Loman NJ. Towards a genomics-informed, real-time, global pathogen surveillance system. Nat Rev Genet. 2018;19:9–20. doi: 10.1038/nrg.2017.88.

7. Quick J, Grubaugh ND, Pullan ST, Claro IM, Smith AD, Gangavarapu K, et al. Multiplex PCR method for MinION and Illumina sequencing of Zika and other virus genomes directly from clinical samples. Nat Protoc. 2017;12:1261–76. doi: 10.1038/nprot.2017.066.

8. Giovanetti M, Faria NR, Lourenço J, Goes de Jesus J, Xavier J, Claro IM, et al. Genomic and Epidemiological Surveillance of Zika Virus in the Amazon Region. Cell Rep. 2020;30:2275–83.e7. doi: 10.1016/j.celrep.2020.01.085.

9. Fauver JR, Petrone ME, Hodcroft EB, Shioda K, Ehrlich HY, Watts AG, et al. Coast-to-Coast Spread of SARS-CoV-2 during the Early Epidemic in the United States. Cell. 2020. doi: 10.1016/j.cell.2020.04.021.

10. Head SR, Komori HK, LaMere SA, Whisenant T, Van Nieuwerburgh F, Salomon DR, et al. Library construction for next-generation sequencing: overviews and challenges. Biotechniques. 2014;56:61–4, 66, 68, passim. doi: 10.2144/000114133.

11. Picelli S, Björklund AK, Reinius B, Sagasser S, Winberg G, Sandberg R. Tn5 transposase and tagmentation procedures for massively scaled sequencing projects. Genome Res. 2014;24:2033–40. doi: 10.1101/gr.177881.114.

12. Bruinsma S, Burgess J, Schlingman D, Czyz A, Morrell N, Ballenger C, et al. Bead-linked transposomes enable a normalization-free workflow for NGS library preparation. BMC Genomics. 2018;19:722. doi: 10.1186/s12864-018-5096-9.

13. Quick J. nCoV-2019 sequencing protocol v1. doi: 10.17504/protocols.io.bbmuik6w.

14. ARTIC Network. artic-ncov2019. accessed May 13, 2020. https://github.com/artic-network/artic-ncov2019. Accessed 13 May 2020.

15. Langmead B, Salzberg SL. Fast gapped-read alignment with Bowtie 2. Nat Methods. 2012;9:357–9. doi: 10.1038/nmeth.1923.

16. ncov2019-artic-nf. Github. https://github.com/connor-lab/ncov2019-artic-nf/tree/f0ba0a1493c9571f4eda161ae8d6afe02d0da570. Accessed 19 May 2020.

17. Li H. Aligning sequence reads, clone sequences and assembly contigs with BWA-MEM. arXiv [q-bio.GN]. 2013. http://arxiv.org/abs/1303.3997.

18. Krueger F. TrimGalore. accessed May 13, 2020. https://github.com/FelixKrueger/TrimGalore. Accessed 13 May 2020.

19. Grubaugh ND, Gangavarapu K, Quick J, Matteson NL, De Jesus JG, Main BJ, et al. An amplicon-based sequencing framework for accurately measuring intrahost virus diversity using PrimalSeq and iVar. Genome Biol. 2019;20:8. doi: 10.1186/s13059-018-1618-7.

20. Li H. seqtk. Github; retrieved May 13, 2020. https://github.com/lh3/seqtk. Accessed 13 May 2020.

21. Quinlan AR, Hall IM. BEDTools: a flexible suite of utilities for comparing genomic features. Bioinformatics. 2010;26:841–2. doi: 10.1093/bioinformatics/btq033.

22. nCoV-2019 Version 3 Amplicon Release. 2020. https://community.artic.network/t/ncov-2019-version-3-amplicon-release/19. Accessed 19 May 2020.

23. Hadfield J, Megill C, Bell SM, Huddleston J, Potter B, Callender C, et al. Nextstrain: real-time tracking of pathogen evolution. Bioinformatics. 2018;34:4121–3. doi: 10.1093/bioinformatics/bty407.

24. Nextstrain - Data Submitter’s FAQ. https://github.com/nextstrain/ncov/blob/master/docs/data_faq.md. Accessed 16 Jun 2020.

25. Shu Y, McCauley J. GISAID: Global initiative on sharing all influenza data - from vision to reality. Euro Surveill. 2017;22. doi: 10.2807/1560-7917.ES.2017.22.13.30494.

26. Batty EM, Kochakarn T, Panthan B, Kumpornsin K, Jiaranai P, Wangwiwatsin A, et al. Genomic surveillance of SARS-CoV-2 in Thailand reveals mixed imported populations, a local lineage expansion and a virus with truncated ORF7a. medRxiv. 2020;:2020.05.22.20108498. doi: 10.1101/2020.05.22.20108498.

27. Artic Network. https://artic.network/quick-guide-to-tiling-amplicon-sequencing-bioinformatics.html. Accessed 19 May 2020.

28. MiSeq performance parameters. https://www.illumina.com/systems/sequencing-platforms/miseq/specifications.html. Accessed 19 May 2020.

