## Supplementary Figures and Tables for "Comparing library preparation methods for SARS-CoV-2 multiplex amplicon sequencing on the Illumina MiSeq platform"

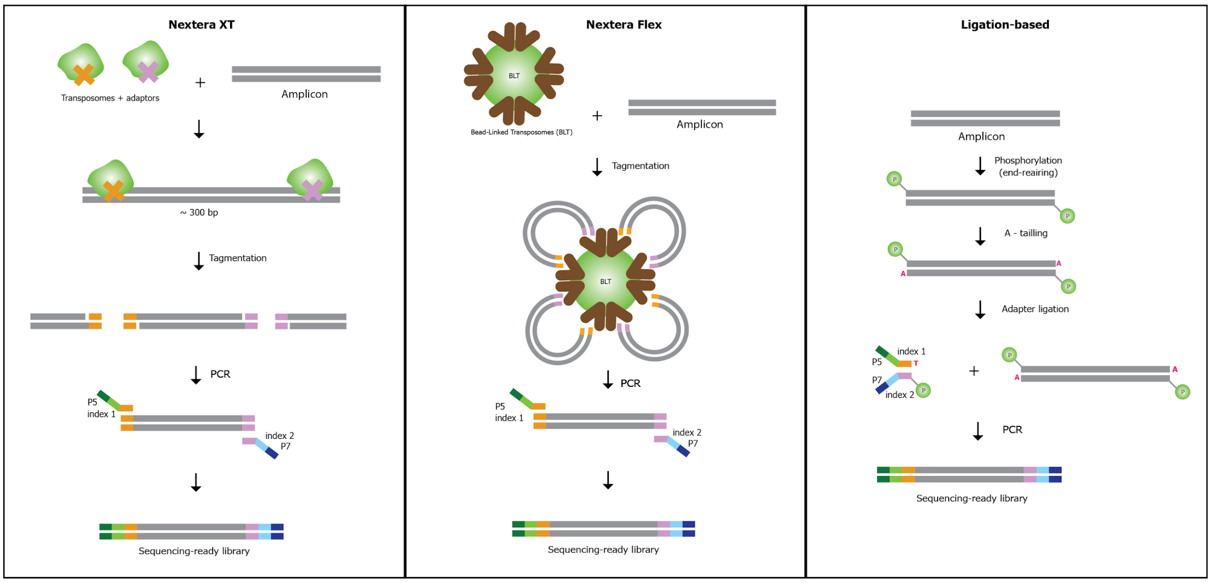


**Supplementary Figure 1.** Schematic workflow for each library preparation method.

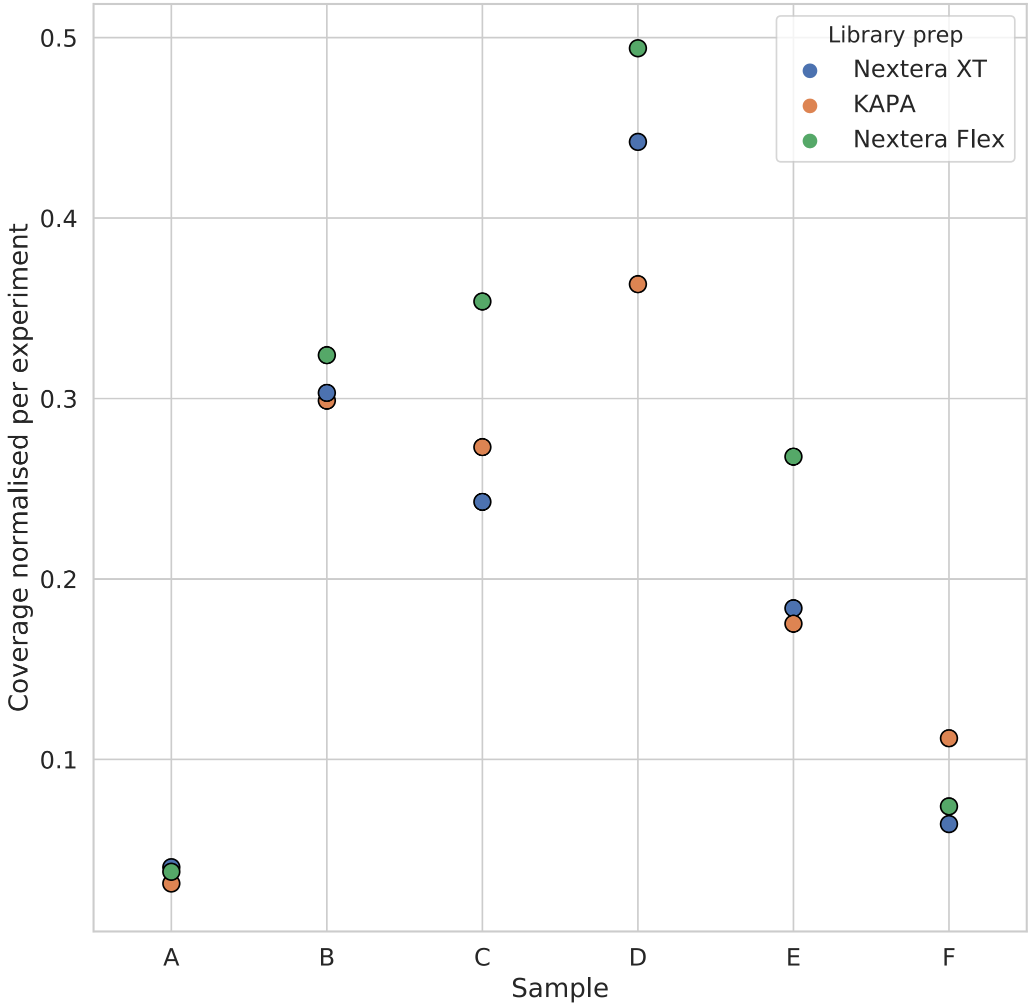


**Supplementary Figure 2.** Median coverage across amplicon 97 for each library. Coverage is normalised within each experiment. Samples A and F have a SNP in the amplicon 97 primer region.

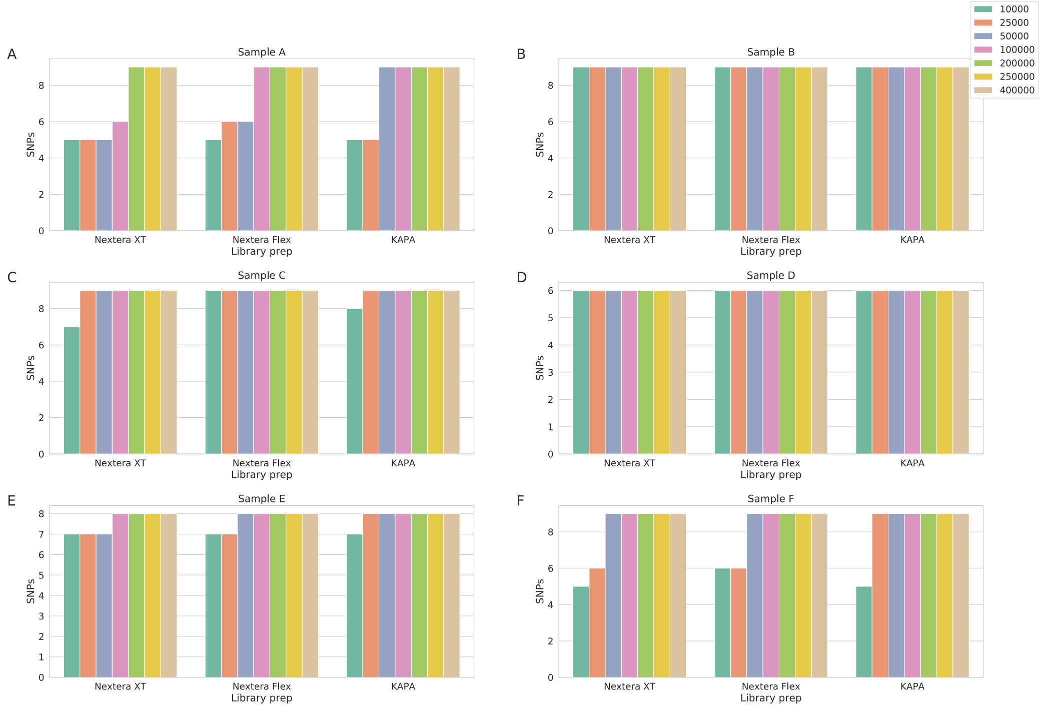


**Supplementary Figure 3.** Number of SNPs called in each sample by library prep method and number of read pairs used.

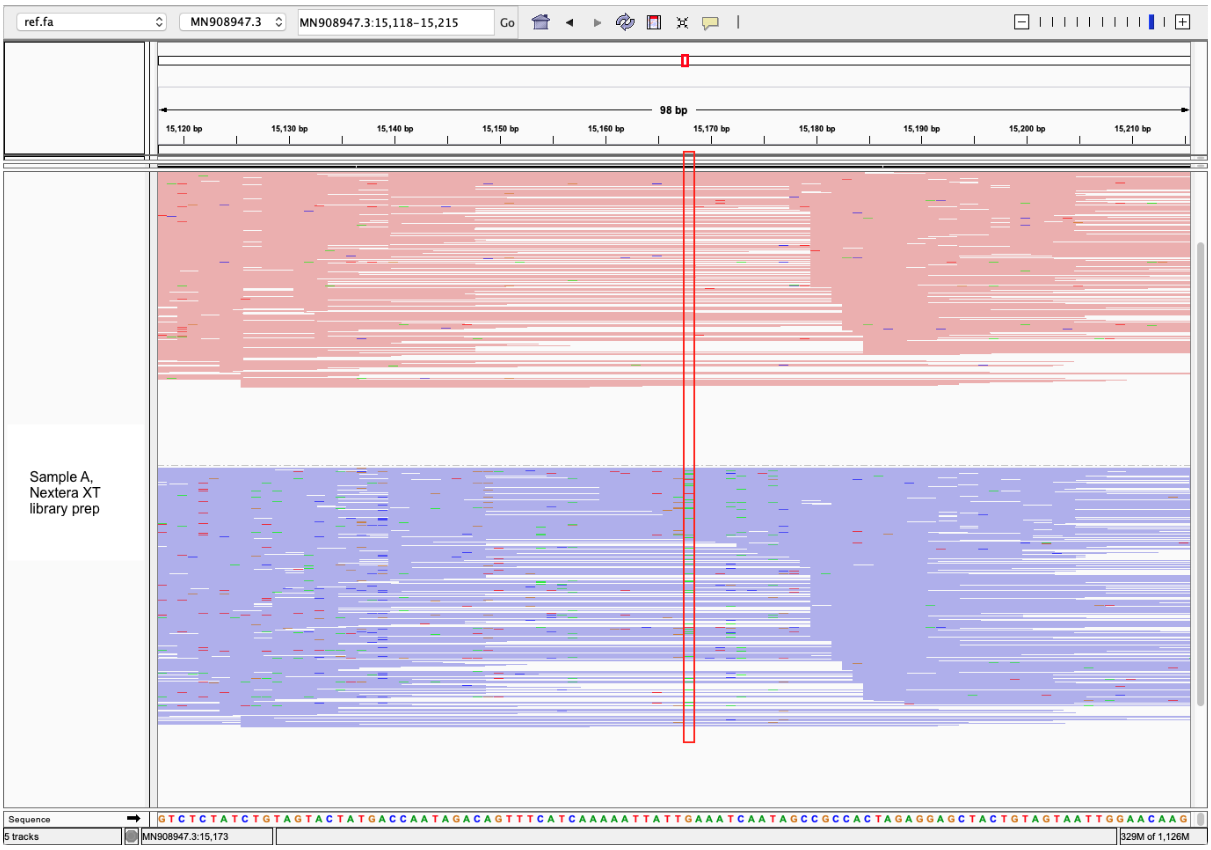


**Supplementary Figure 4**. Screenshot of IGV alignment around position 15168 in the Nextera XT library prep sample, showing the variant position in a red box. The alternative base call is seen only in negative strand reads.

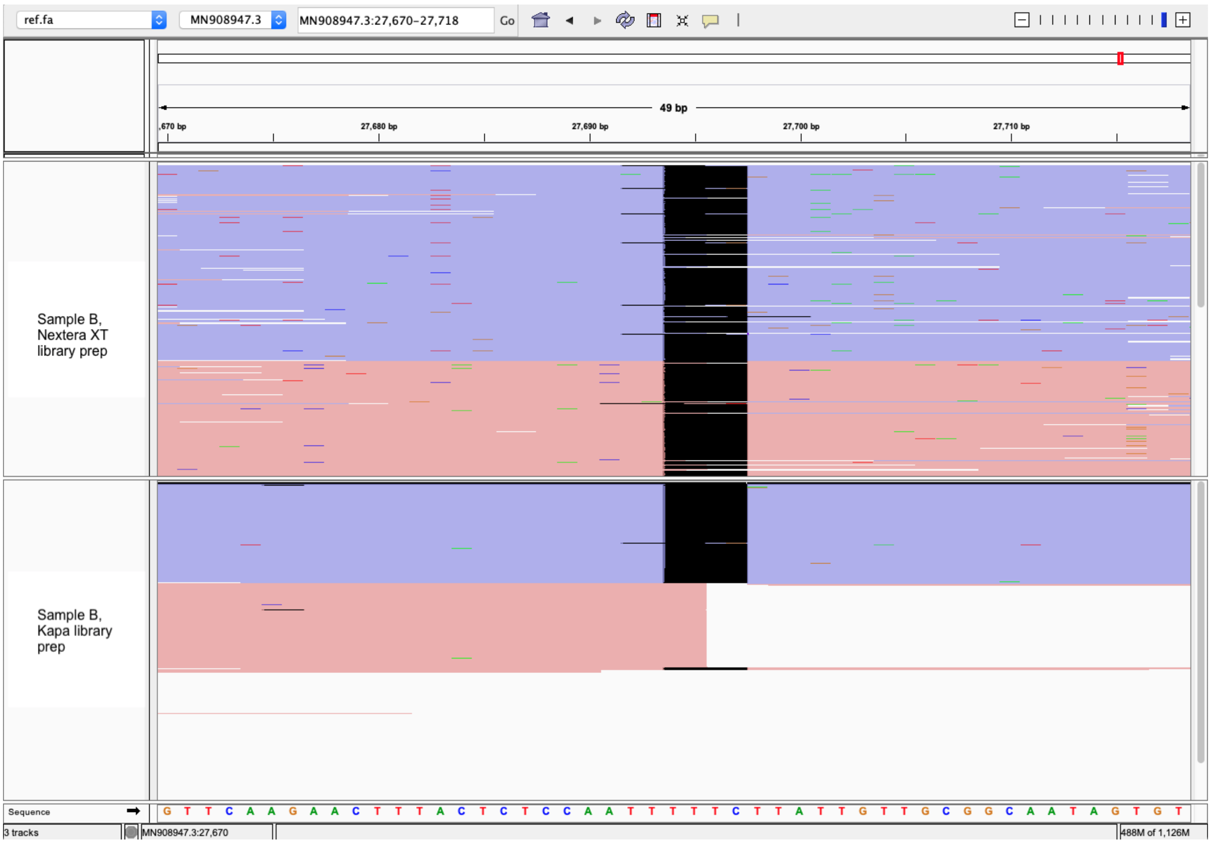


**Supplementary Figure 5.** Screenshot of IGV alignment around position 27693 in the Nextera XT and KAPA library prep for the same sample. The black bar shows the deletion, which is not seen in the positive strand reads in the KAPA library prep (pink bars), as the read alignment is soft-clipped.

### Supplementary Tables

| Sample | $C_t$ (Orf1a) | $C_t$ (RdRp) |
|---|---|---|
| A | 22 | 22.4 |
| B | 13.5 | 13.6 |
| C | 27.3 | undetermined |
| D | 24.1 | 22.1 |
| E | 20.6 | 20.6 |
| F | 18.59 | 19 |

**Supplementary Table 1**. $C_t$ values of RNA samples used in this study. The qRT-PCR was carried out previously by Ramathibodi Hospital for clinical diagnostic test.

| Sample | Amplicon yield (ng) | | Library concentration (ng/µl) | | | Library yield (ng) | | | Library fragment size (bp) | | |
|---|---|---|---|---|---|---|---|---|---|---|---|
| | Pool 1 | Pool2 | Nextera XT | Nextera Flex | Ligation-based | Nextera XT | Nextera Flex | Ligation-based | Nextera XT | Nextera Flex | Ligation-based |
| A | 1137 | 1296 | 5.08 | 2.15 | 30.2 | 254 | 64.5 | 906 | 308 | 465 | 512 |
| B | 594 | 867 | 5.41 | 2.06 | 58.2 | 270.5 | 61.8 | 1746 | 259 | 467 | 538 |
| C | 477 | 651 | 5.61 | 2.11 | 27 | 280.5 | 63.3 | 810 | 260 | 467 | 512 |
| D | 762 | 1071 | 5 | 2.32 | 42.8 | 250 | 69.6 | 1284 | 269 | 466 | 518 |
| E | 221.7 | 312 | 6.51 | 2.57 | 36.4 | 325.5 | 77.1 | 1092 | 272 | 469 | 530 |
| F | 519 | 477 | 6.27 | 2.77 | 81.1 | 313.5 | 83.1 | 2433 | 262 | 470 | 556 |
| NTC1 | 18 | 17.28 | 7.2 | 0.987 | 0.216 | 360 | 29.61 | 6.48 | 290 | 459 | 512 |
| NTC2 | 20.73 | 17.64 | 6.28 | 0.662 | 1.87 | 314 | 19.86 | 56.1 | 320 | 440 | 512 |

**Supplementary Table 2**. RT-PCR and library preparation QC. Concentrations were measured using Qubit fluorometer. Library fragment sizes were based on Fragment Analyzer System. NTC, negative control.

| Sample | Library prep | Reads mapped to SARS-CoV-2 | Reads remaining after Ivar quality filtering and primer trimming |
|---|---|---|---|
| NTC1 | Nextera XT | 3165 | 464 |
| NTC2 | Nextera XT | 2251 | 310 |
| NTC1 | Nextera Flex | 7602 | 3787 |
| NTC2 | Nextera Flex | 4873 | 2341 |
| NTC1 | KAPA | 20358 | 268 |
| NTC2 | KAPA | 5907 | 779 |

**Supplementary Table 3**. The number of reads which were mapped to SARS-CoV-2 after the initial filtering step of our pipeline, and the number which remained after final quality filtering and trimming. NTC, negative control.
